## Supplementary Figures and Tables for "Integrating taxonomic, functional, and strain-level profiling of diverse microbial communities with bioBakery 3"

### Supplementary Figures

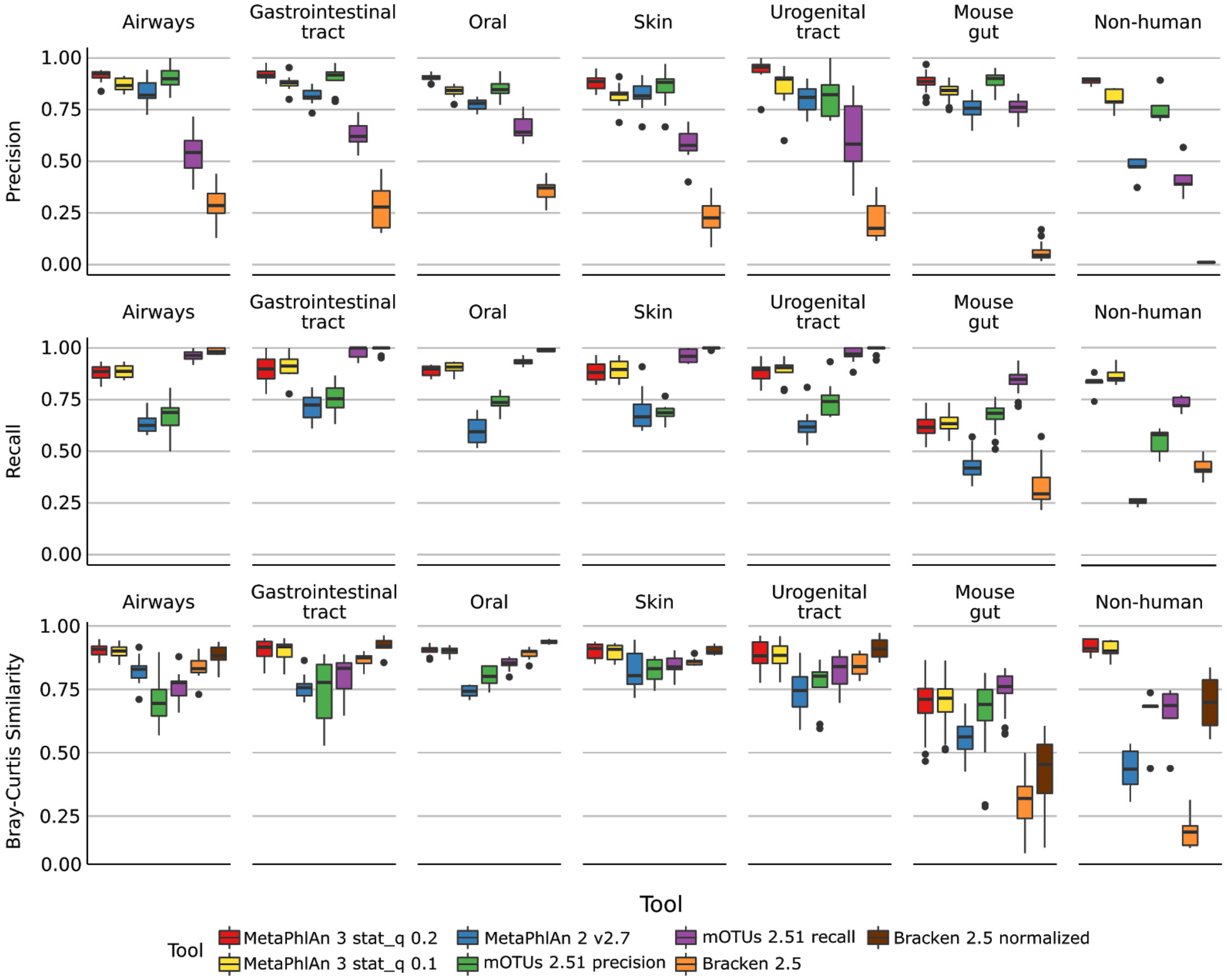


**Supplementary Figure 1:** Performance metrics (Precision, Recall, Bray-Curtis similarity) of MetaPhlAn 3.0, MetaPhlAn2, mOTU, and Bracken species-level profiling of the CAMI human-associated, CAMI mouse gut, and non-human datasets. Bray-Curtis similarity index is calculated on arcsine-square-root transformed relative abundances

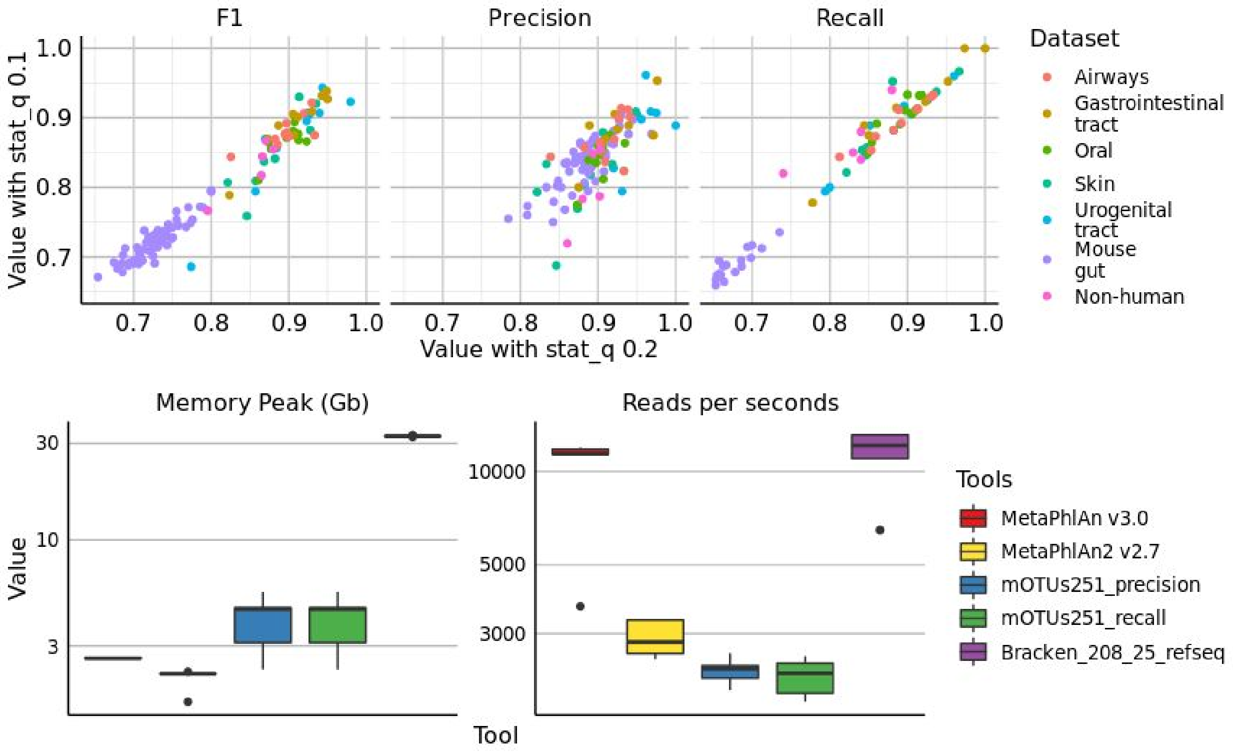


**Supplementary Figure 2: (top)** Scatter plots of precision, recall, and F1 score, of all the synthetic metagenomes profiled with MetaPhlAn 3 using stat_q=0.2 (default value for MetaPhlAn 3) and stat_q=0.1 (rho = 0.97). **(bottom)** Comparison of memory usage (maxRSS) and speed of taxonomic profilers included in the evaluation. Each tool was run on 5 HMP metagenomes using 1 thread.

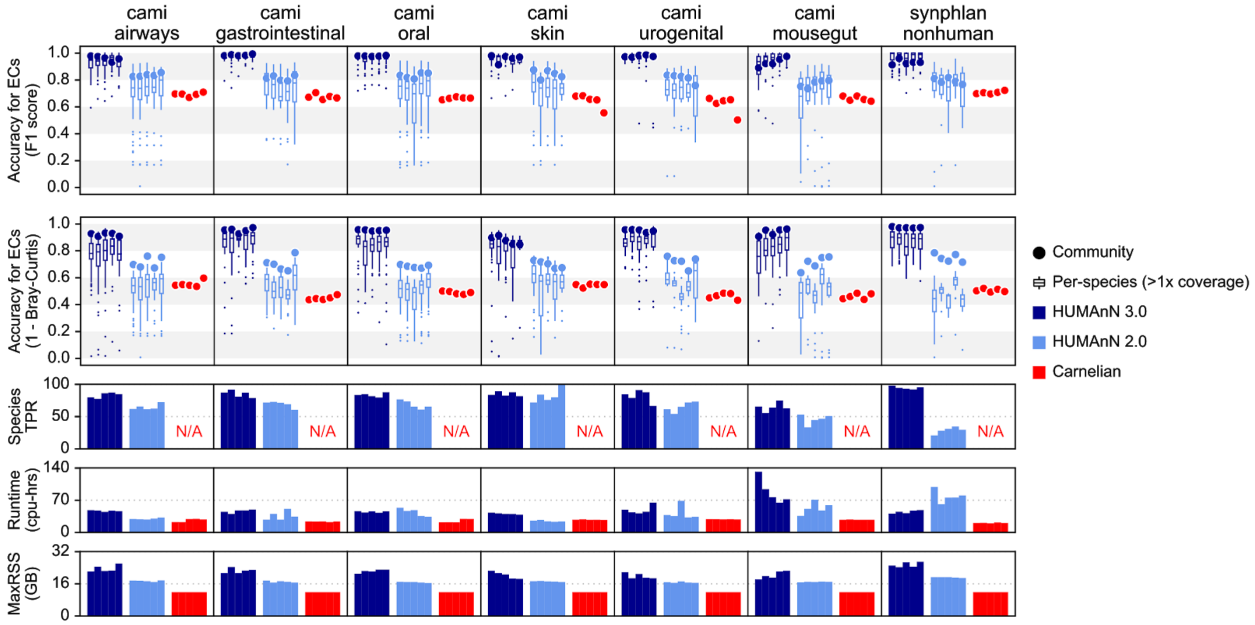


**Supplementary Figure 3:** This figure expands Fig. 1D from the main text to further compare HUMAnN 3, HUMAnN 2, and Carnelian on the basis of F1 score for accuracy of enzyme commission (EC) family detection, runtime (cpu-hrs), and peak memory usage (MaxRSS).

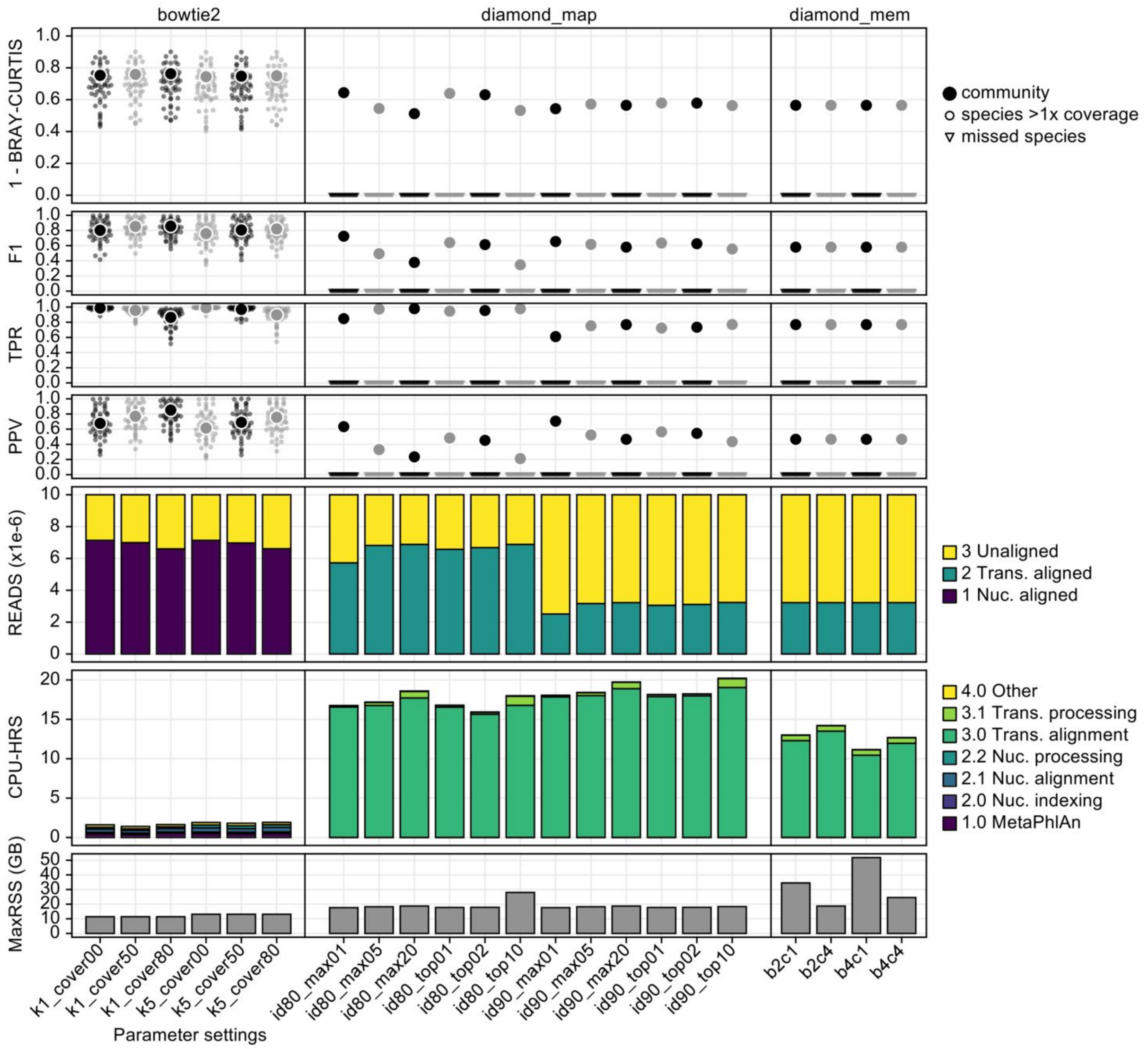


**Supplementary Figure 4:** This figure summarizes our initial optimization of HUMAnN 3 based on the synphlan-humanoid metagenome with a UniRef90 gold standard. Pangenome search (bowtie2 phase) was evaluated in "--bypass-translated-search" mode and translated search (diamond phase) was evaluated in "--bypass-nucleotide-search" mode. Left ("bowtie2") column: We compared accuracy and performance requesting 1 vs. 5 hits from Bowtie 2 and performing post hoc filtering of target sequences requiring 0% (i.e. no filtering), 50%, and 80% of sites to be hit. HUMAnN 3 defaults to a single hit (unchanged from HUMAnN 2) but requires 50% coverage of database sequences (similar to HUMAnN 2's translated search filter). Center ("diamond_map") column: We compared a variety of DIAMOND stringency filters during translated search. HUMAnN 3 uses alignments with >80% identity within 1% score of the best alignment (id80_top01), which is more sensitive (but otherwise similar) to the HUMAnN 2 default (id90_max20). Right column ("diamond_mem"): We evaluated different memory utilization settings in DIAMOND, but kept the DIAMOND default (b2c4) in both HUMAnN 2 and 3.

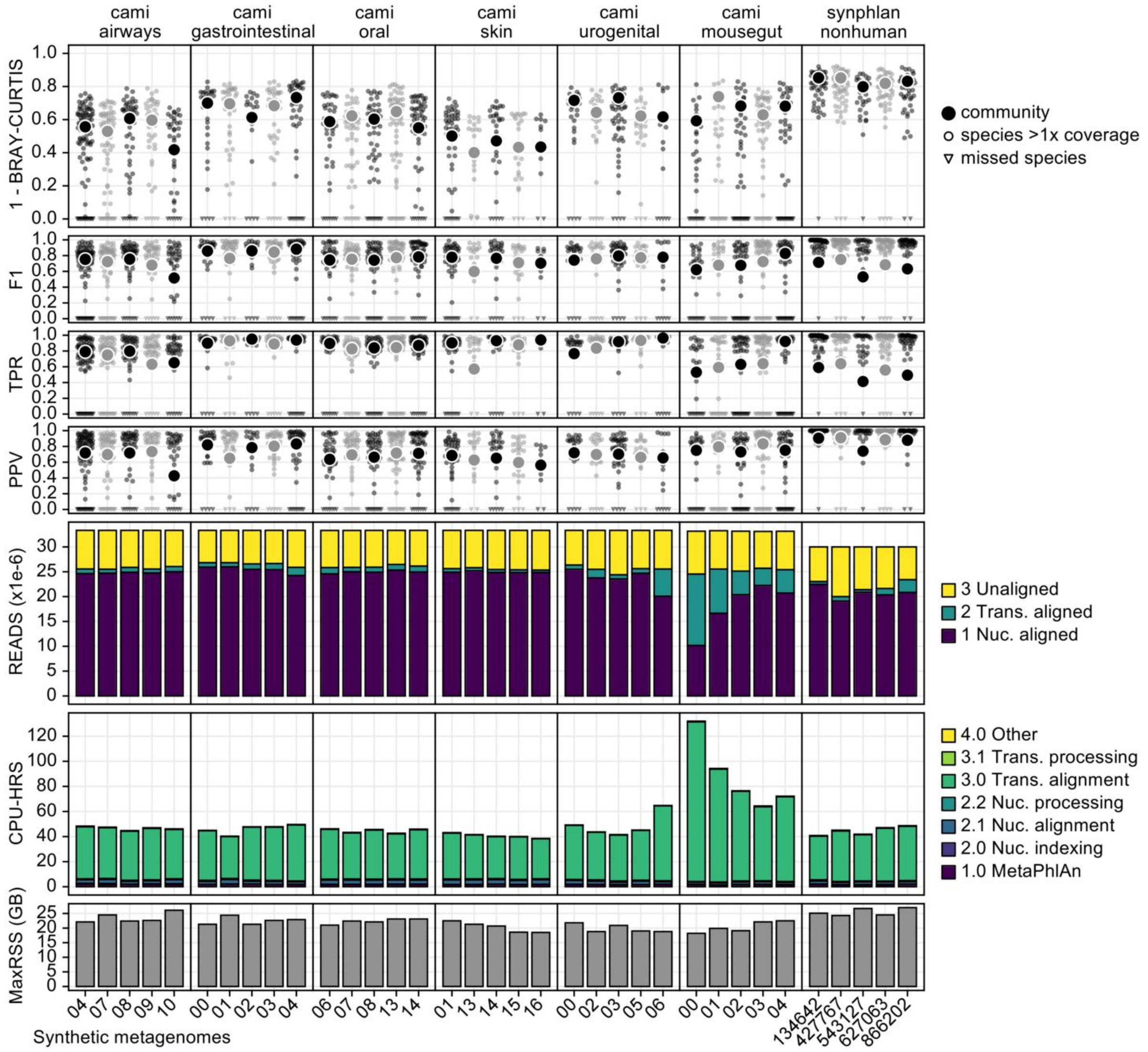


**Supplementary Figure 5:** This figure provides a high-resolution view of HUMAnN 3's performance in the evaluations of main-text Fig. 1D (accuracy and performance on CAMI and non-human-associated metagenomes). The top four rows (1 - BC, F1, TPR, and PPV) detail measures of accuracy for UniRef90-level protein families at the community (large dot) and well-covered-species (small dots) levels. The "READS" row indicates the stage of HUMAnN 3's tiered search where sample reads were aligned; ~75% of most samples' reads were explained, with the vast majority of the reads assigned by known pangenomes outside of the CAMI mousegut samples (which relied more heavily on translated search for explanations). The "CPU-HRS" row indicates the time spent in various phases of HUMAnN 3's tiered search, with the translated search step dominating overall runtime. The MaxRSS row indicates the peak memory usage (in GBs) for each sample, and was consistently in the 20-25 GB range.

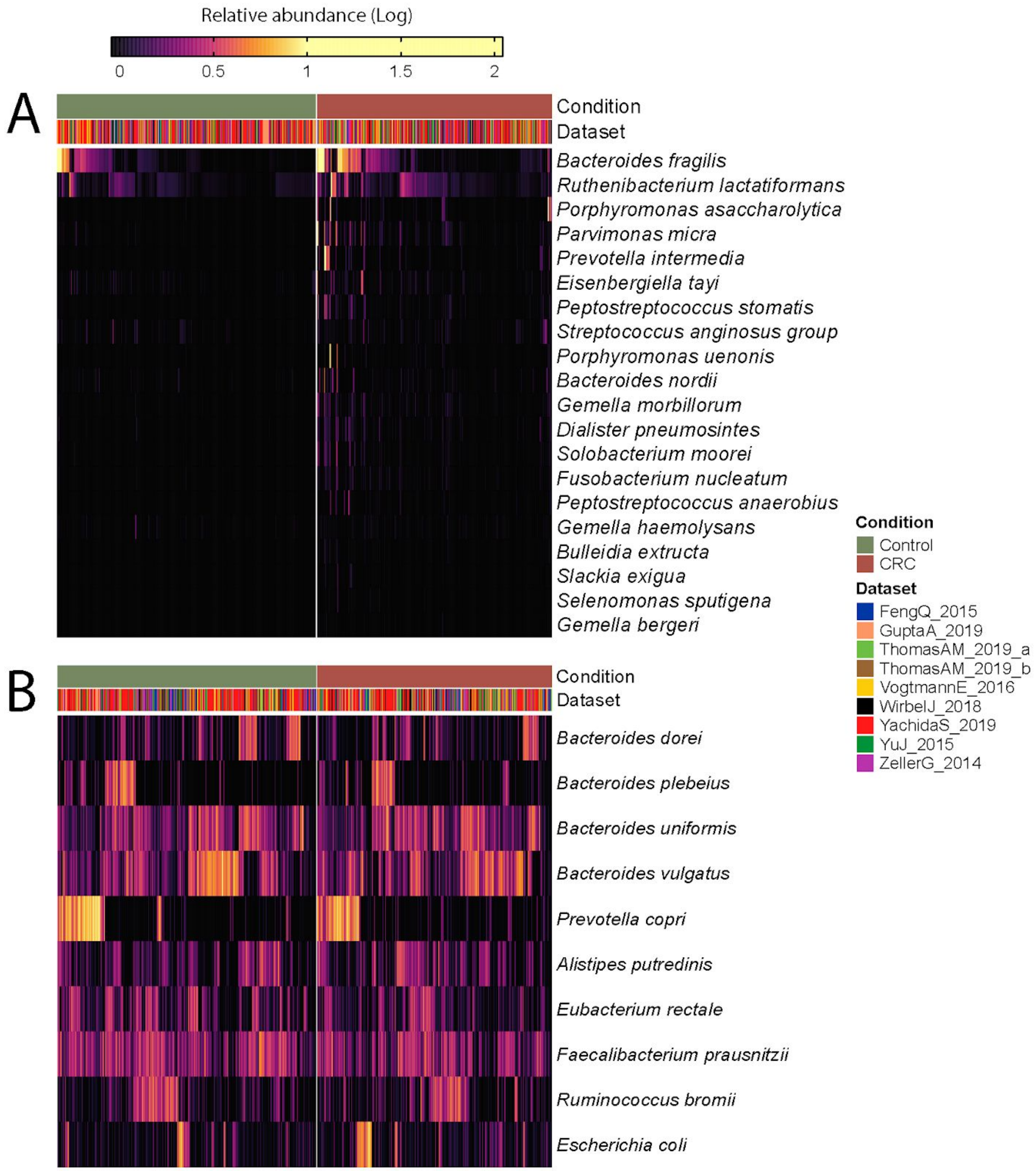


**Supplementary Figure 6:** Log-transformed relative abundances of the top 20 MetaPhlAn 3 species associated with colorectal cancer **(A)** and top 10 most abundant species **(B)** identified with a meta-analysis on 1,262 samples.

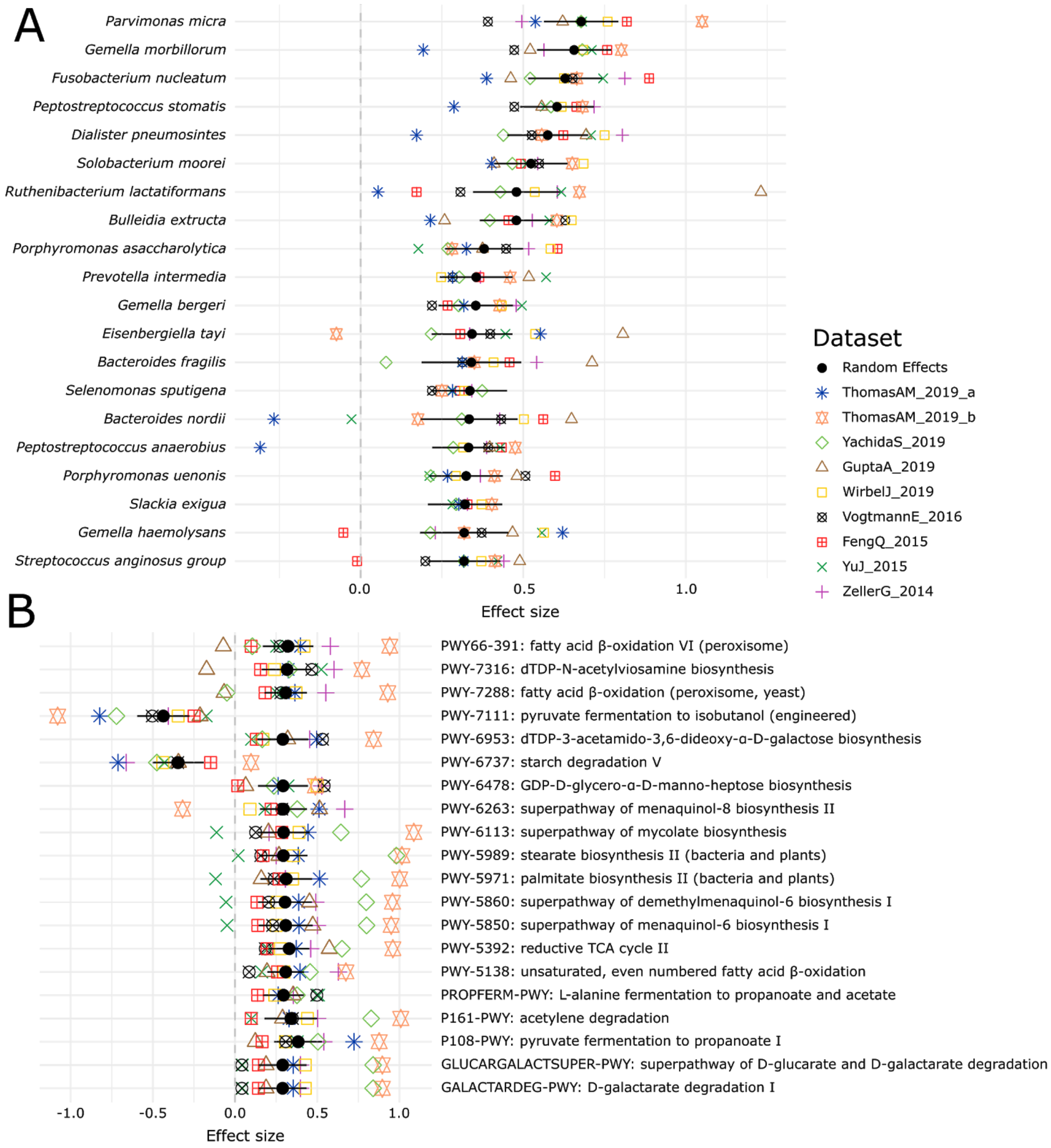


**Supplementary Figure 7:** Meta-analysis of the CRC datasets on the MetaPhlAn 3.0 species-level relative abundances **(A)** and relative abundance of MetaCyc pathway profiles generated with HUMAnN 3 **(B)**.

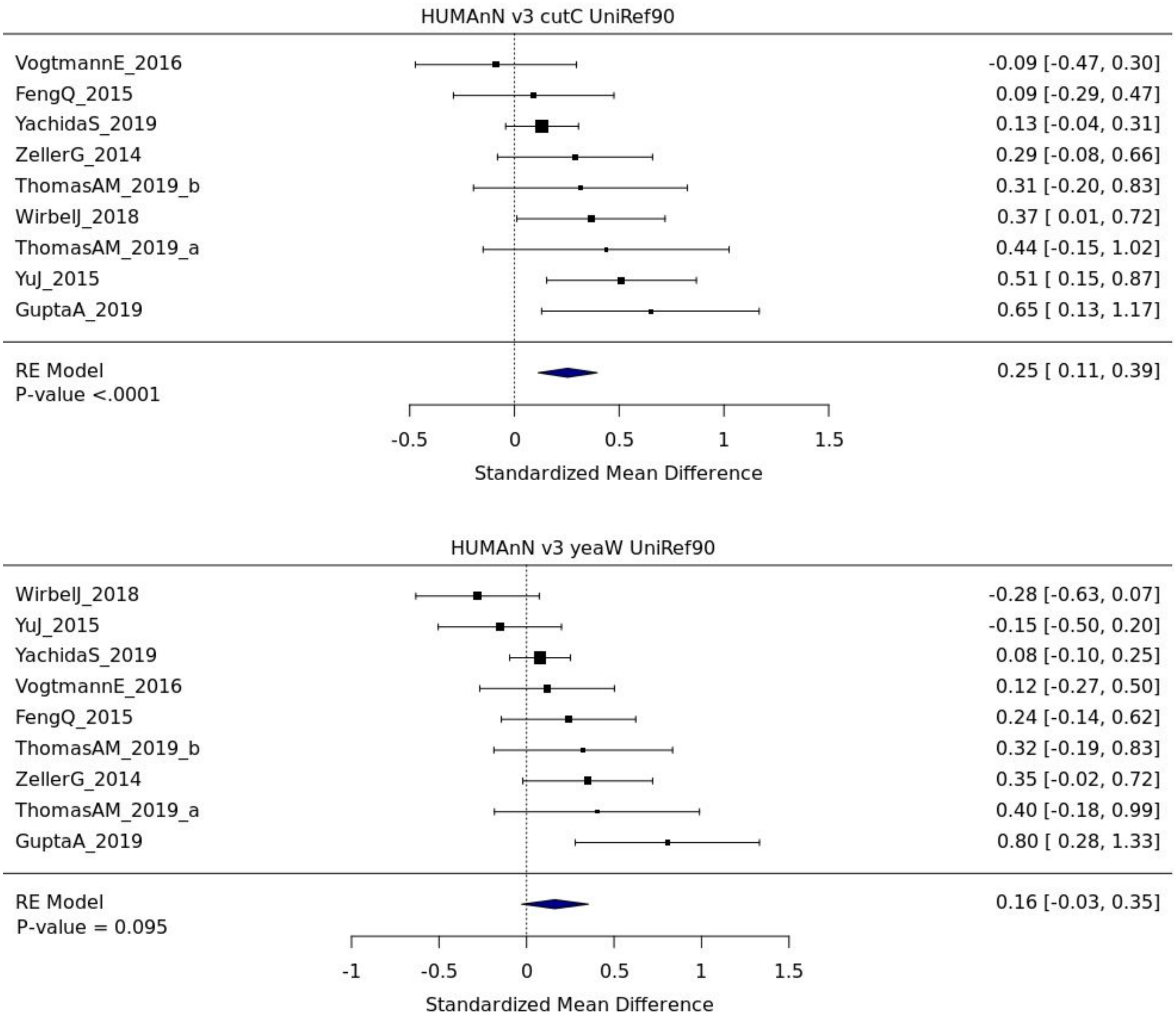


**Supplementary Figure 8:** Forest plot reporting effect sizes calculated using a meta-analysis of standardized mean differences and a random effects model on cutC and yeaW relative abundances between CRC and control samples.

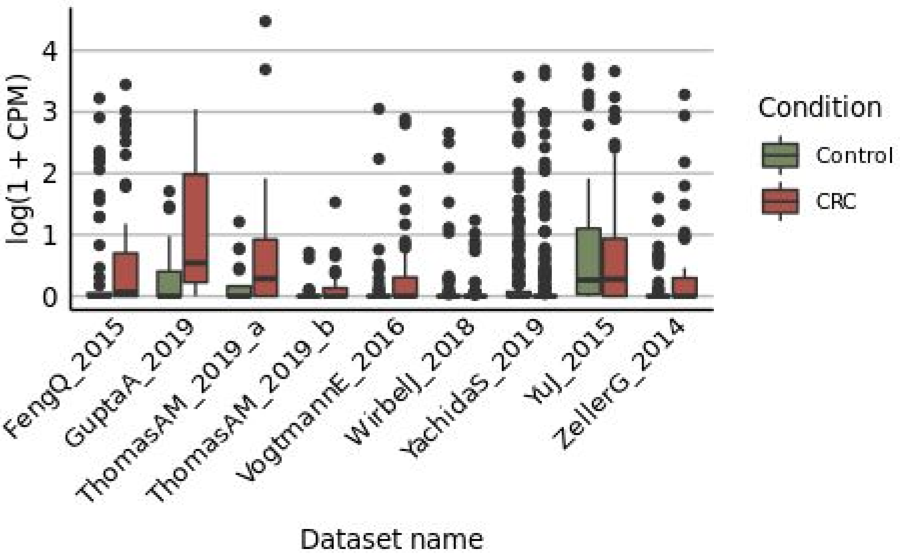


**Supplementary Figure 9:** Distribution of yeaW gene relative abundance (log10 count-per-million normalized) extracted from HUMAnN gene family profiles.

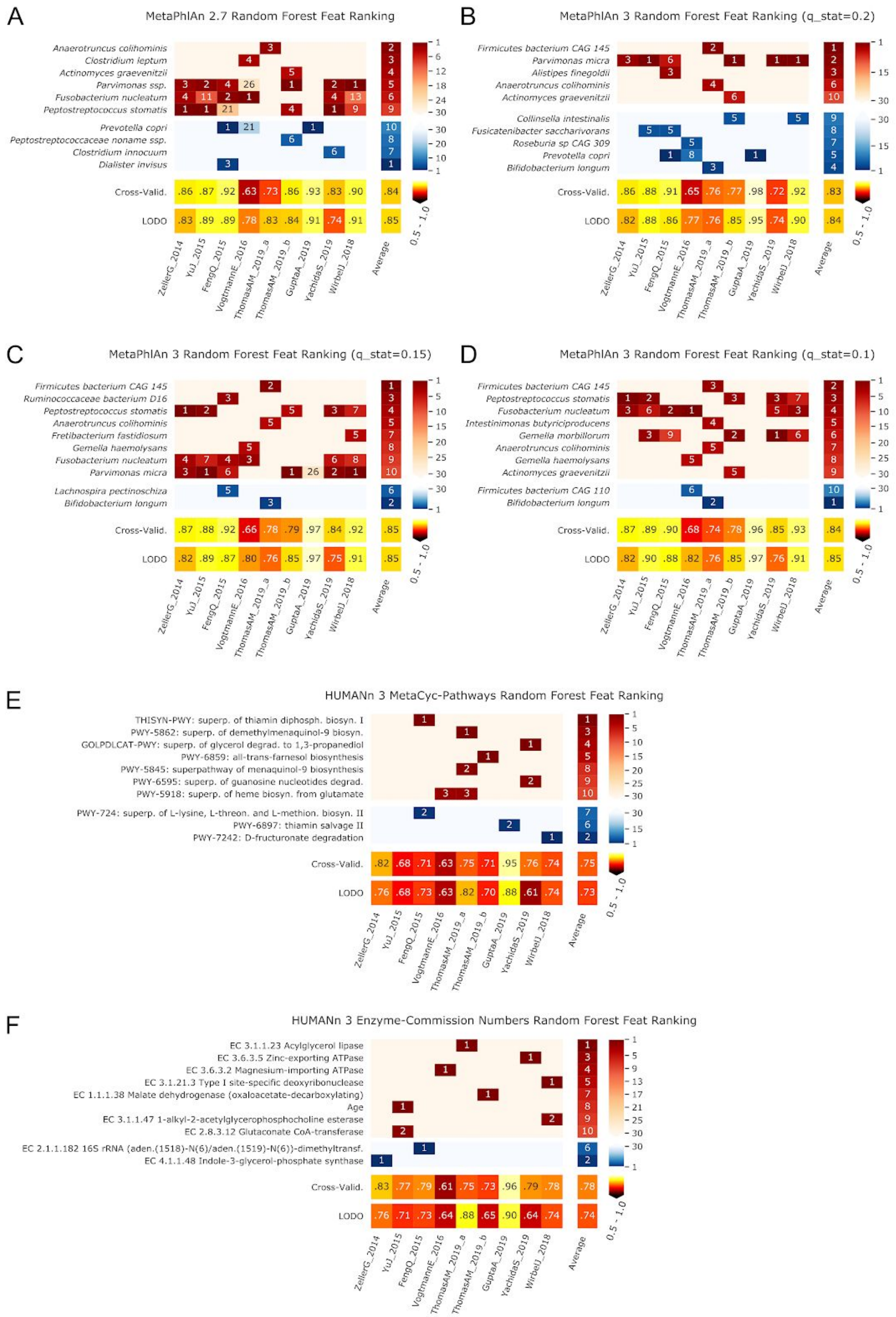


**Supplementary Figure 10:** Features identified by the random-forest analysis on the species profiled with MetaPhlAn2 and MetaPhlAn 3 using different values of q_stat, and by HUMANn 3 grouping UniRef90 in MetaCyc pathways and Enzyme Commission numbers.

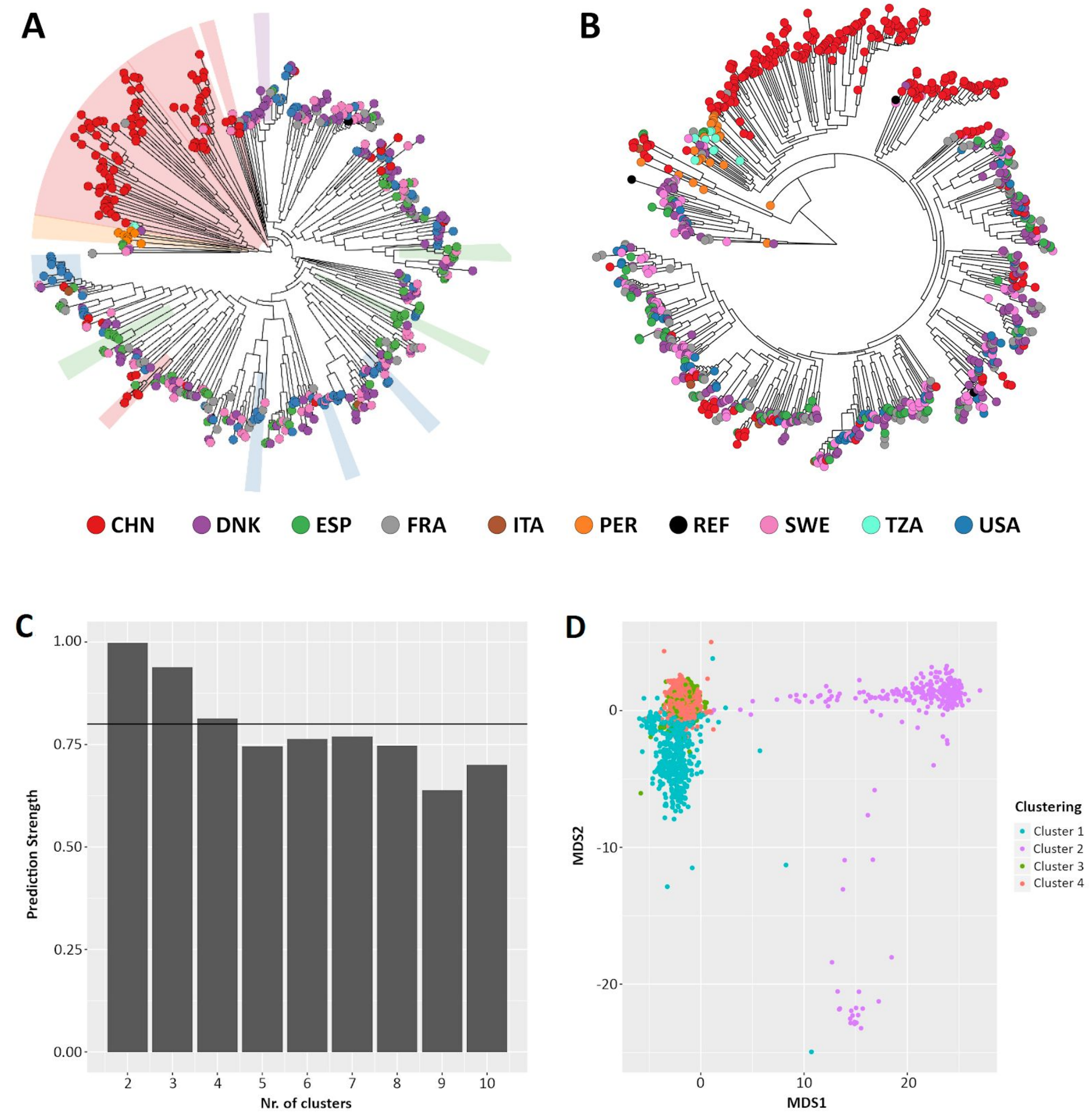


**Supplementary Figure 11:** Comparison between StrainPhlAn **(A)** and StrainPhlAn 3 **(B)** strain level profiling capabilities. *Ruminococcus bromii* species was profiled on 1,590 metagenomes. **(C)** Prediction strength at different cluster numbers and **(D)** PAM clustering results on the StrainPhlAn 3 phylogenetic distance matrix expose four optimal clusters of *Ruminococcus bromii* strains.

### Supplementary Tables

**Supplementary Table 1: Average values of F1 scores of MetaPhlAn 3, MetaPhlAn2, mOTUs2, and Kraken species-level profiles computed on the 123 synthetic metagenomes.**

| Tool | Airways | Gastrointestinal tract | Oral | Skin | Urogenital tract | Mouse gut | Non-human |
|---|---|---|---|---|---|---|---|
| **MetaPhlAn v3.0 stat_q 0.2** | 0.880 | 0.894 | 0.869 | 0.853 | 0.869 | 0.722 | 0.830 |
| **MetaPhlAn v3.0 stat_q 0.1** | 0.896 | 0.908 | 0.898 | 0.883 | 0.906 | 0.728 | 0.855 |
| **MetaPhlAn2 v2.7** | 0.723 | 0.761 | 0.672 | 0.751 | 0.701 | 0.544 | 0.330 |
| **mOTUs251_precision** | 0.768 | 0.824 | 0.787 | 0.761 | 0.779 | 0.770 | 0.632 |
| **mOTUs251_recall** | 0.683 | 0.768 | 0.772 | 0.720 | 0.727 | 0.800 | 0.529 |
| **Bracken_208_25_refseq** | 0.440 | 0.426 | 0.525 | 0.359 | 0.339 | 0.091 | 0.021 |

**Supplementary Table 2: bioBakery 3 software improvements.**
https://www.dropbox.com/s/sbar6dggsrohz1m/Supplementary_table_2_biobakery_comparision.xlsx?dl=0

**Supplementary Table 3: Mean and ranked values of Bray-Curtis dissimilarity and arcsine-square-root normalized Bray-Curtis dissimilarity obtained by MetaPhlAn 3, MetaPhlAn2, mOTUs2, and Kraken on the synthetic metagenomes considered in the evaluation.**
https://www.dropbox.com/s/ptcmml1mkri002x/Supplementary_table_3_taxonomic_profiling_bray_curtis.xlsx?dl=0

**Supplementary Table 4: Comparison of runtime and memory consumption of MetaPhlAn 3, MetaPhlAn2, mOTUs2, and Kraken+Bracken on the 5 HMP metagenomes.**

| Tool | Elapsed time (mean h) | Elapsed time (sd h) | Memory Peak (mean Gb) | Memory Peak (sd Gb) | Reads per second (mean) | Reads per second (sd) |
|---|---|---|---|---|---|---|
| **MetaPhlAn v3.0** | 3.1120 | 2.65 | 2.614 | 0.01 | 10031.2 | 3,560.25 |
| **MetaPhlAn2 v2.7** | 8.2183 | 3.58 | 2.079 | 0.27 | 2911.2 | 400.08 |
| **mOTUs251_precision** | 10.6094 | 4.92 | 4.034 | 1.30 | 2283 | 234.26 |
| **mOTUs251_recall** | 11.5189 | 6.18 | 4.035 | 1.30 | 2186 | 310.86 |
| **Bracken_208_25_refseq** | 2.3504 | 1.40 | 32.529 | 0.27 | 11163 | 2,759.93 |

**Supplementary Table 5: MetaPhlAn 3 taxonomic profiles and HUMAnN 3 functional profiles of the 1,262 CRC samples.**
https://www.dropbox.com/s/3jn5tgdx7ssw5v2/Supplementary_table_5_CRC_metaphlan_humann_profiles.xlsx?dl=0

**Supplementary Table 6: MetaPhlAn 3 species-level and HUMAnN 3 pathway abundances CRC meta-analysis results.**
https://www.dropbox.com/s/s4zlop6jz2f35ek/Supplementary_table_6_CRC_metaanalysis_metaphlan_results.xlsx?dl=0

**Supplementary Table 7: MetaPhlAn 3 species merged according to the species-level genome bin (SGB) system.**
https://www.dropbox.com/s/r2fv71jxw81y6h4/Supplementary_table_7_metaphlan3_merged_species_with_SGB.xlsx?dl=0

**Supplementary Table 8: Number of distinct MetaPhlAn 3 markers per species.**
https://www.dropbox.com/s/yl7638cps97i72f/Supplementary_table_8_metaphlan3_markers_per_species.xlsx?dl=0

**Supplementary Table 9: Per-sample OPAL binary measures (true positive, false positive, false negative, precision, recall, F1 score) computed on MetaPhlAn 3, MetaPhlAn2, mOTUs2, and Kraken species-level profiles computed on the 123 synthetic metagenomes.**
https://www.dropbox.com/s/eokxfegceb6b7t4/Supplementary_table_9_evaluation.xlsx?dl=0

**Supplementary Table 10: Metadata of all the 1,262 samples from the 10 CRC datasets.**
https://www.dropbox.com/s/kwvdzcjoni0kug7/Supplementary_table_10_CRC_metaanalysis_datasets.xlsx?dl=0
